# Evaluating trait-based sets for taxonomic enrichment analysis applied to human microbiome data sets

**DOI:** 10.1101/2022.05.16.492155

**Authors:** Quang P. Nguyen, Anne G. Hoen, H. Robert Frost

## Abstract

**Background:** Set-based pathway analysis is a powerful tool that allows researchers to summarize complex genomic variables in the form of biologically interpretable sets. Since the microbiome is characterized by a high degree of inter-individual variability in taxonomic compositions, applying enrichment methods using functionally driven taxon sets can increase both the reproducibility and interpretability of microbiome association studies. However, there is still an open question of which knowledge base to utilize for set construction. Here, we evaluate microbial trait databases, which aggregate experimentally determined microbial phenotypes, as a potential avenue for meaningful construction of taxon sets.

**Method:** Using publicly available microbiome sequencing data sets (both 16S rRNA gene metabarcoding and whole-genome metagenomics), we assessed these trait-based sets on two criteria: first, do they cover the diversity of microbes obtained from a typical data set, and second, do they confer additional predictive power on disease prediction tasks when assessed against measured pathway abundances and PICRUSt2 prediction.

**Results:** Trait annotations are well annotated to a small number but most abundant taxa within the community, concordant with the concept of the core-peripheral microbiome. This pattern is consistent across all categories of traits and body-sites for whole genome sequencing data, but much more heterogenous and inconsistent in 16S rRNA metabarcoding data due to difficulties in assigning species-level traits to genus. However, trait-set features are well predictive of disease outcomes compared against predicted and measured pathway abundances. Most important trait-set features are more interpreable and reveal interesting insights on the relationship between microbiome, its function, and health outcomes.

## Introduction

Advancements in high-throughout sequencing technologies have allowed researchers to characterize the identity and functional potential of a large proportion of microorganisms in human-associated microbiomes. This has enabled efficient study of the link between health outcomes and the microbiota without reliance on currently limited culture-based approaches [35]. As such, there has been an increase in microbiome profiling studies, primarily aiming towards identifying specific microbes that are differentially abundant between groups of individuals defined by an exposure or disease state vs a control population [71]. However, such analyses face unique computational and statistical challenges [36], which includes addressing the burden of multiple testing and providing meaningful biological interpretations.

This challenge of understanding the results of microbiome analyses in the broader context of biological systems mirrors that of other high-throughput data sets. One approach that has proven to be fruitful in human genomic studies is gene set testing (or pathway analysis) which focuses on analyzing the coordinated expression of groups of genes (termed gene sets or pathways) [41]. From a statistical perspective, set-based statistics are are more reproducible and have greater power compared to their gene-level counter parts [22]. The true benefit of set-based approaches, however, is the ability to incorporate *a priori* knowledge of specific cellular processes [38]. Microbiome differential abundance analyses can also benefit from set-based approaches instead of a microbe-centric approach. In addition to statistical benefits such as reduced dimensionality and sparsity [33, 45], set-based approaches are also more reflective of the underlying biology. Like genes, microbes act in concert with co-abundant partners to drive biochemical processes that interact with the host, thereby impacting health outcomes [66]. For example, when comparing patients with inflammatory bowel disease against healthy subjects, microbes thought to be disease-causing for inflammatory bowel disease were also strongly co-occurring [21], suggesting that they might jointly contribute to the microbiome-disease causal pathway instead of acting as independent factors. This is also represented in the development of therapies, where products often contain multiple strains of bacteria [3, 18]. Furthermore, organizing microbes into functionally-driven groups (also termed “guilds” [66]) is also congruent with the perspective that human microbiomes are complex ecosystems whose properties emerge from localized interactions between microbial communities representing individuals that exploit and contribute to their environment in similar ways [19].

Unfortunately, there is currently limited research in curating and evaluating appropriate microbe annotations similar to the transcriptomic literature. Repositories like the Molecular Signatures Database (MSigDB) [38] aggregate information about gene function across multiple sources, incorporating both laboratory results and computational inferences. Even though similar databases such as Disbiome [29] and MSEA [33] exist, they are usually human-centric and define microbial groups based on their potential to be pathological rather than through common biochemical roles. As such, these databases are limited in generating meaningful hypotheses linking taxonomic changes to ecosystem function especially in novel disease conditions. Trait-based analysis [4, 40, 63], with its long history in traditional macroecological studies [19, 34], is a promising approach to address this gap. Traits directly represent microbial physiological characteristics and metabolic phenotypes (for example, sulfur reduction, nitrate utilization, or gram positivity) and therefore can serve as annotations for potential ecosystem function. For 16S rRNA gene sequencing data sets, where one can only obtain taxonomic abundances, performing enrichment analysis on trait-based sets can elucidate the taxa-function relationship and identify microbial processes that are differentially active between healthy and diseased patients. For whole genome metagenomic data sets, traits still offer unique perspectives. First, traits are often sourced from the long history of laboratory experiments such as journal articles and *Bergey’s Manual of Systematic Archaea and Bacteria* [57] which is different from homology-based sequence queries typically performed to profile gene family abundances. Second, traits are complex phenotypes that represent multiple molecular pathways, which means that they are more comparable to higher-order pathway annotations in hierarchical databases such as Kyoto Encyclopedia of Genes and Genomes (KEGG) [32] and MetaCyc [12]. As such, utilizing traits as the source to group microbes into functional and phenotypical categories can assist in interpreting microbiome profiling studies, and generating mechanistically meaningful hypotheses that link ecosystem function and its taxa.

Even though trait-based approaches have been utilized in various studies [4, 23, 34, 63], to our knowledge there is currently no effort to formalize trait-based databases in terms of microbial sets and evaluate their utility in a typical enrichment analysis of 16s rRNA metabarcoding or metagenomic data. Here, we constructed taxon sets from pre-existing trait databases at both the species and genus level. Then, we computed the coverage of these traits across different human-associated environments and sequencing approaches. Finally, we evaluated whether trait-based set features confer predictive capacity for diseased individuals compared to measured (from whole genome sequencing data) and predicted (from PICRUSt2 [17]) pathway abundances. Finally, we identified the most important features for prediction and assessed whether they matched existing literature on the microbiome-disease relationship of interest.

## Material and methods

All analyses were performed in the R programming language (version 4.1.2) [52] and the Python programming language (version 3.10.4). All graphics were generated using `ggplot2`, `ggsci`, `patchwork`. Tabular data manipulation was performed using `pandas` for python, and `tidyverse` suite of packages for R. Additional packages utilized include: `BiocSet`, `taxizedb`, `phyloseq`, `TreeSummarizedExperiment`. For enrichment analyses, we leveraged the CBEA [45] method (version 1.0.1) developed previously by our lab. All analyses were performed using the `snakemake` workflow [43]. All reproducible code and intermediate analysis products can be found on GitHub (qpmnguyen/microbe set trait).

### Generating taxonomic sets from trait databases

We utilized pre-compiled trait databases from previous publications: Madin et al. 2020 [40] and Weissman et al. 2021 [63]. The former was chosen due to the fact that it is the most comprehensive compilation of microbial (bacteria and archaea) physiological traits based on existing sources to date. The latter is a newer database that hand curates traits specifically for human microbiomes based on Bergey’s manual. Both of these databases source their trait assignments primarily from biochemical and microbiological laboratory experiments over genomic-based annotation. We focused our analyses on categorical traits, namely metabolism, gram stain, enzymatic pathways, sporulation, motility, cellular shape, and substrate utilization. We are particularly interested in traits belonging to the class of enzymatic pathways and substrate utilization as they represent functions that most directly impact the microbe-host relationship [60].

We combined both databases into a joint knowledge base and constructed sets for each available categorical trait. Additionally for the Madin et al. database, we updated data entries sourced from Genomes Online Database (GOLD) [44] due to the fact that compared to other compiled sources, GOLD is continuously updated via community submissions. We grouped all traits belonging to the same National Center for Biotechnology Information (NCBI) species-level identifier. When there are conflicts in assigning traits, we prioritized Weissman et al. over Madin et al. and GOLD due to its hand curated nature. If there are ambiguities in taxonomic assignment in the Weissman et al. source, we considered that trait to be missing. The exceptions to the above logic are enzymatic pathways and substrate utilization categories where trait values across sources for the same species are concatenated instead of reconciled. For example, if a species A has entries from multiple databases suggesting the presence of “nitrogen degredation” and “ammonia degredation”, then instead of attempting to chose the best annotation based on source we assumed that species A has the capacity to metabolize both nitrogen and ammonia.

All traits are defined at the species level via NCBI identifiers, however, due to restrictions for 16S rRNA gene sequencing data sets to resolve beyond the genus level [30], we also assigned traits to each genus based on a two-step process for each major trait category:

- A hypergeometric test is used to ascertain whether the genus is underrepresented in the database based on the total number of species assigned to that genus in NCBI Taxonomy [55] compared to our trait database. If a genus is underrepresented in our database (i.e. the proportion of species number of genera in the database is significantly less than what one would expect if one were to randomly draw species from the NCBI database), then the trait is not assigned to that genus since we do not have enough information. Specifically, we assessed *P* (*X*≤ *x*) at *α* = 0.05 where *X* ∼ *Hypergeometric*(*k, N, K*), with *x* as the total number of species assigned to that genus in the database with an assigned value for the trait category of interest, *k* as the total number of species in the database with an assigned value for the trait category, *N* as the total number of species in NCBI Taxonomy, and *K* as the total number of species assigned to the genus in NCBI Taxonomy.
- For all genera that are well represented in the database, we then assessed the proportion of species under that genus that have the trait. If over 95% of species of a given genus have the trait, then the trait is assigned to the genus.

We then defined trait-based sets using the aforementioned assignments. Each trait value with a category, e.g. “obligate anaerobic” from the category “metabolism”, is defined as a set with elements representing the species (or genus) annotated to that trait value. In the analysis stage, each identified taxon within a data set is matched to a trait based on their NCBI identifier. For 16S rRNA gene metabarcoding data sets, we matched all amplicon sequence variants (ASV) with traits belonging to the genus level NCBI identifier matched to the ASV sequence. All processed databases and resulting taxonomic sets can be found on GitHub in the analysis repository.

### Evaluation data sets

We evaluated trait-based sets on publicly available 16S rRNA gene metabarcoding and whole-genome metagenomic data sets. For study-specific metabarcoding data sets, we obtained data directly from associated European Nucleotide Archive (ENA) repositories and re-processed raw sequence files into ASV tables using the `dada2` QIIME 2 (version 2022.2) plugin [7, 10]. Taxonomic classification was performed using a pre-trained weighted naive bayes model [6, 31] using the SILVA NR 99 database version 138 [51] available via QIIME 2. For all our metagenomic data sets, we downloaded taxonomic and pathway abundance tables directly from the `curatedMetagenomicData` R package [47] (2021-10-19 snapshot), which processed the data via the `bioBakery` [2] metagenomic data processing pipeline by the package authors. Data from the Human Microbiome Project (HMP) was obtained using the `HMP16SData` [54] (for metabarcoding data) and `curatedMetagenomicData` [47] (for metagenomic data) R packages.

To assess trait annotation coverage, we utilized data from both Phase I and II of the HMP [16] as it contains surveys for multiple human-associated environments from healthy subjects. For predictive and concordance analyses, we focused on colorectal cancer (CRC) and inflammatory bowel disease (IBD) as study conditions. Both CRC and IBD are well represented across both metabarcoding and metagenomic data sets, allowing comparisons across sequencing methodology. Furthermore, these conditions are also under active study within the microbiome literature, which improves the ability to interpret the biological significance of the results. For CRC, we utilized data from Zeller et al. [70], Feng et al. [20], Gupta et al. [24], Hannigan et al. [26], Thomas et al. [56], Vogtmann et al. [61], Wirbel et al. [64], Yachida et al. [68], and Yu et al. [69]. For IBD, we utilized data from the integrative HMP [50], Gevers et al. [21], Hall et al. [25], Ijaz et al. [28], Li et al. [37], Nielsen et al. [46], and Vich Vila et al. [59]. A detailed description of each data set and data-processing procedures is available in the Supplementary Materials.

### Coverage analysis

In this analysis, we sought to identify how well trait databases cover the taxonomic diversity of different human-associated environments. We leveraged healthy samples from multiple body sites from Phase I and II of the HMP [16]. We quantified coverage as a per-sample measure considering both taxa absence/presence and its abundance.

- For each sample, we computed the proportion of taxa that is present (non-zero counts) assigned to at least one trait (a sample-level trait-specific richness).
- For each sample, we computed the proportion of reads assigned to taxa that is present and annotated to at least one trait (a sample-level trait-specific evenness).

In addition to coverage stratified by trait categories and body sites, we also generated category-specific and site-specific coverage values by averaging across all sites or categories, respectively.

### Prediction analysis

We also aimed to evaluate whether trait-based features can add information for microbiome-based disease prediction compared to other data inputs. Here, we generated sample-level enrichment scores for each trait using `CBEA` and utilized them as inputs to a standard random forest model [8]. Model fitting was done using `scikit-learn` [48] where all parameters were set to default values with the exception of the total number of trees per ensemble (500) and the total number of features considered per split (equal to the square root of the total number of features). We compared model performance using trait enrichment scores against measured and PICRUSt2 predicted pathway abundances (for metagenomic and metabarcoding data sets, respectively).

Model performance was measured using the area under the receiver operating characteristic curve (AUROC) and Brier scores [9]. These metrics and associated confidence intervals were obtained by fitting and evaluating the model via a 10-fold cross-validation procedure. To obtain calibrated predictive probabilities for Brier scores, we applied Platt’s method (using `CalibratedClassifierCV`) with 5-fold cross validation nested within the training fold and used the ensemble model to generate test set probabilities [49].

In order to identify which features are important to the disease prediction process, for each input type we re-split the entire data set into train/test splits (80% training data). We then refitted our calibrated random forest model on the training set as described above. Since our final model is an ensemble of calibrated random forest classifiers, we obtained feature importance values as the average across all calibrated cross validation folds (*N* = 10). Feature importance per random forest model is defined based on the implementation in `scikit-learn` as the decrease in Gini impurity when the feature is split averaged across all decision trees in a forest.

## Results

### Trait-based taxonomic sets Database coverage

We computed the coverage for each trait category across each body site in the HMP data set. Fig 1 illustrates results for species-level trait assignment for samples profiled via whole genome metagenomics. In panel A, coverage is evaluated as the total number of taxa present per sample annotated to a trait (a measure of cross-trait richness), while in panel B coverage is the total number of reads assigned to taxa annotated to a trait (a measure of cross-trait evenness). Richness provides a general overview on how many members of a community is assigned to a trait, while evenness accounts for their relative abundances by up-weighting species that have high abundance across all samples.

**Figure 1.**
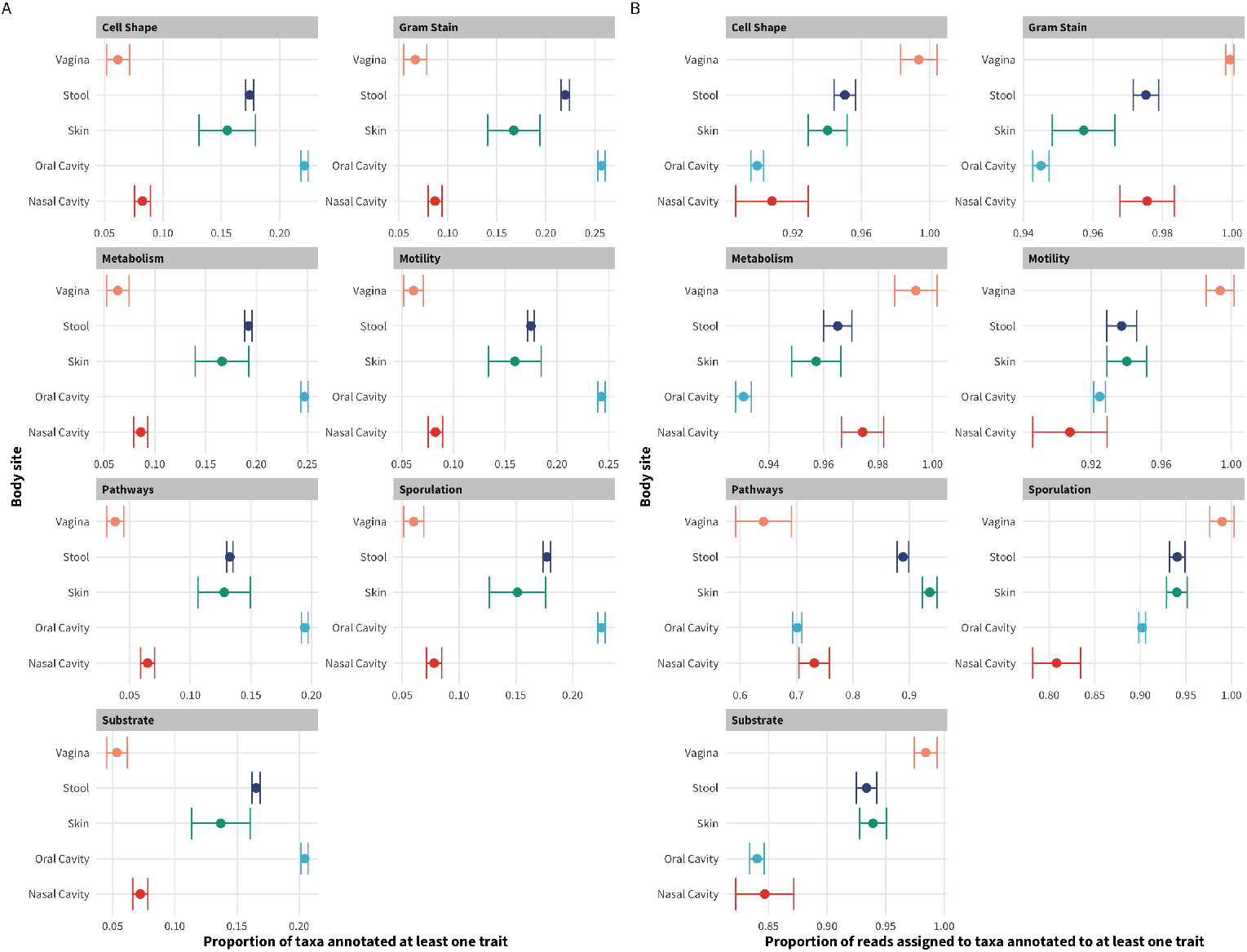
Trait annotation coverage across different body sites for the HMP data set profiled using whole genome shotgun sequencing. Panel **(A)** illustrates the proportion of present taxa per sample annotated to at least one trait. Panel **(B)** illustrates the proportion of reads assigned to taxa annotated to at least one trait which accounts for taxa relative abundances. Each plot facet represents different trait categories that were evaluated. Error bar represents the standard error of the evaluation statistic of interest across the total number of samples evaluated per body site.

Overall, for any body site, at most 25% of taxa are assigned to a trait, but when considering the proportion of reads, coverage increased to more than 80%. This shows that traits are usually well annotated to the most abundant taxa. This pattern holds for samples profiled with 16S rRNA gene sequencing (Fig S1), even though the proportions were much lower due to difficulties in aggregating species level traits to genus. For many body sites and trait category combinations, traits could not be assigned to any taxa.

We also observed heterogeneity in the annotation coverage across different body sites and trait categories. For richness, nasal cavity and vaginal body sites were the lowest in coverage, with less than 5% of taxa annotated with at least one trait across all trait categories while conversely, oral cavity sites consistently had the highest coverage under this metric. This pattern was reversed when considering coverage as the proportion of assigned reads per sample, but overall values were consistently high. Averaging coverage across body sites (Fig 2) also supports this observation, showing overall that the proportion of reads covered are similar across all body sites despite differences in the proportion of present taxa covered by trait annotations. Similar results were observed for sites profiled with 16S rRNA gene sequencing (Fig S1), where oral sub-sites have the highest coverage across both richness and evenness metrics but, on average, all sites were similar in coverage statistics. Surprisingly, stool samples were low in coverage across multiple categories despite being one of the well studied systems.

**Figure 2.**
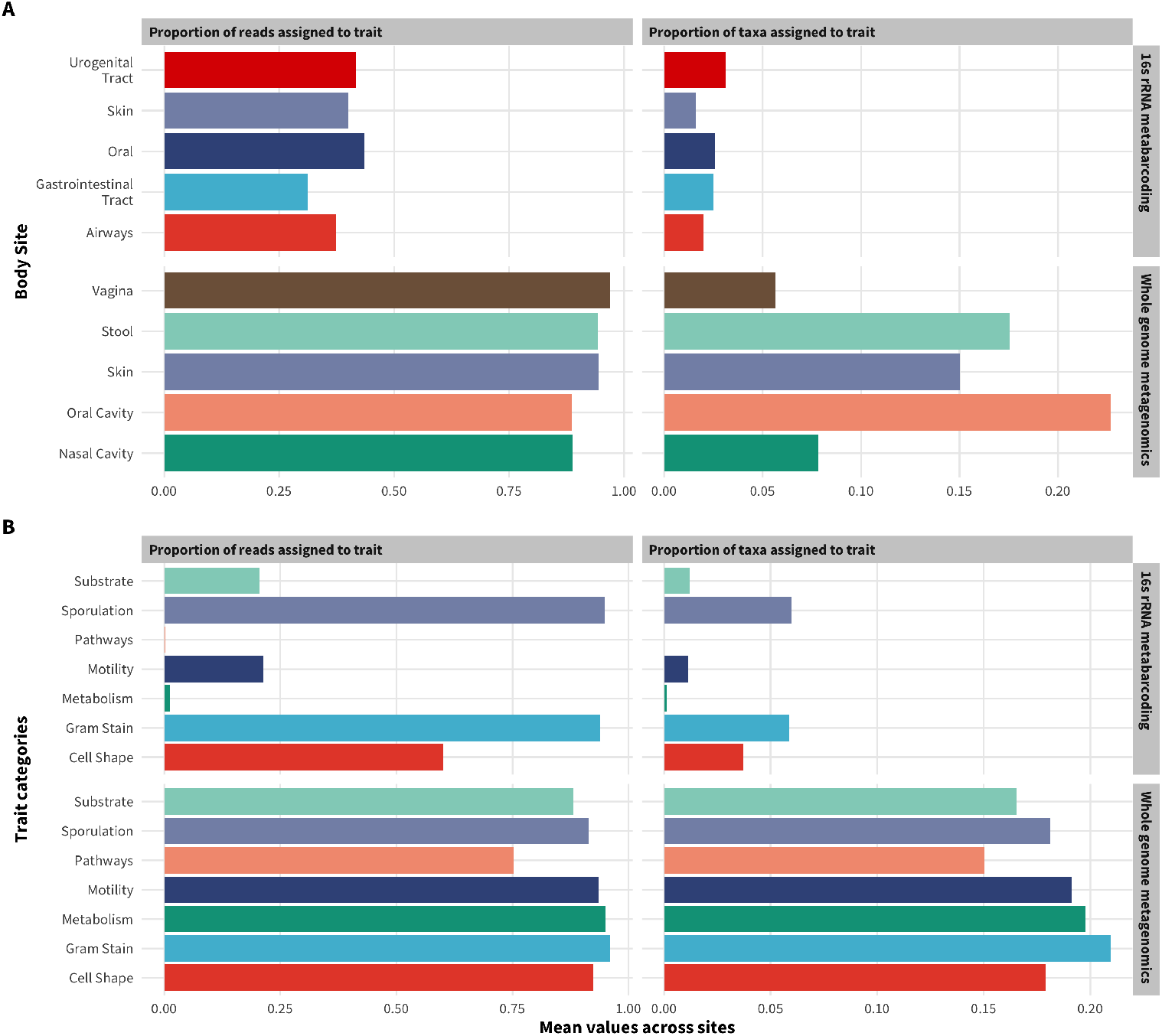
Trait annotation coverage statistics for HMP data across samples profiled with both 16S rRNA gene metabarcoding and whole genome metagenomics. Panel **(A)** illustrates coverage statistics for each body site averaged across evaluated samples and trait categories. Panel **(B)** illustrates statistics for each trait category averaged across evaluated samples and body sites.

We also stratified our coverage analyses by trait categories (Fig 1, Fig 2, Fig S1). For samples profiled with whole genome sequencing, all trait categories are evenly covered, with about 15% - 20% of taxa were annotated to a trait of any category. However, these taxa comprise around 75% to 100% of the total reads per sample suggesting that the overall read level coverage is very high. However, in samples profiled with 16S rRNA gene sequencing, the overall coverage value across categories is low.Sporulation, substrate utilization and motility are the most covered category while pathways and metabolism has no coverage at all.

### Predictive analysis

To determine the utility of trait-based sets, we generated enrichment scores for covered traits using CBEA [45] for evaluated data sets and compared the predictive performance of using trait-set enrichment scores as inputs compared to alternative functional-based predictors. We evaluated two disease conditions, CRC and IBD, with data sets drawn from both 16S rRNA gene metabarcoding and whole genome metagenomic profiling techniques. We fitted a calibrated random forest model to each input type and computed predictive performance as AUROC (discriminatory power) and Brier scores (probability estimates) using 10-fold cross-validation.

For the CRC prediction task, traits covered 2.7% of taxa and 27.3% of reads for the 16S rRNA gene metabarcoding data set, while for the whole genome sequencing data set, traits covered 9.1% of taxa and 87.2% of reads. For the IBD prediction task, traits covered 1.61% of taxa and 26.7% of reads for 16S rRNA gene metabarcoding data set, while for the whole genome sequencing data set, traits covered 6.6% of taxa and 91.2% of reads.

Fig 3 illustrates results of our model evaluations. Overall, enrichment scores for trait-sets are as good as other alternate function-based predictors at discriminating between case and control patients across both CRC and IBD conditions. Aside from pure discrimination power, models fitted on CBEA trait-set scores are also equivalent in approximating predicted probabilities. This is surprising especially for the 16S rRNA gene metabarcoding data sets, where the trait coverage is low. Even though the differences in performance is not significant, there are instances where trait-set scores perform slightly better than their pathway abundance counterparts. Since trait-features are also more descriptive, utilizing them can increase interpretation while also not scarificing performance.

**Figure 3.**
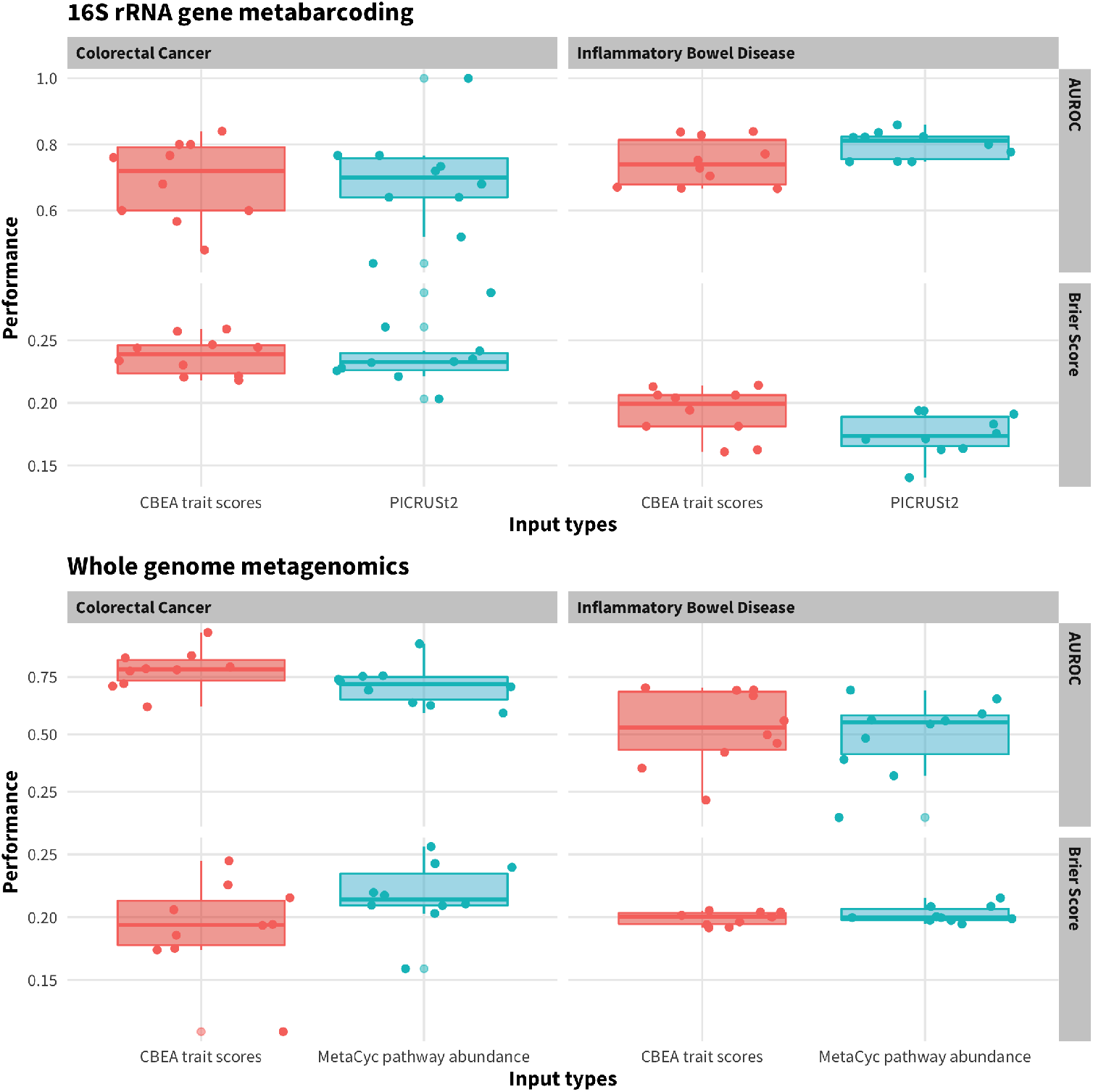
Predictive performance of a calibrated random forest model across different disease conditions, profiling technique, and performance metrics. Scores were obtained via 10-fold nested cross validation where within each training fold there is a 10-fold cross validation procedure to calibrate predicted probabilities. For each condition and data type, CBEA trait-set scores were compared against MetaCyc pathway abundances from relevant sources (measured abundances for whole genome sequencing data sets and PICRUSt2 predicted abundances for 16S rRNA gene metabarcoding data sets)

In addition to predicted performance, we also identified the top 10 features that are most important for model fitting. Since our model involves a 10-fold cross-validation procedure within the training set to calibrate predicted probabilities, top features are identified using the mean feature importance value across the 10 folds. Fig 4 illustrates results for whole genome metagenomic data sets while Fig S2 illustrates results for the 16S rRNA gene metabarcoding data sets. Even though these are the top 10 features, the observed mean feature importance statistics are low, suggesting that no individual features were definitively the most important in discriminating between patient classes.

**Figure 4.**
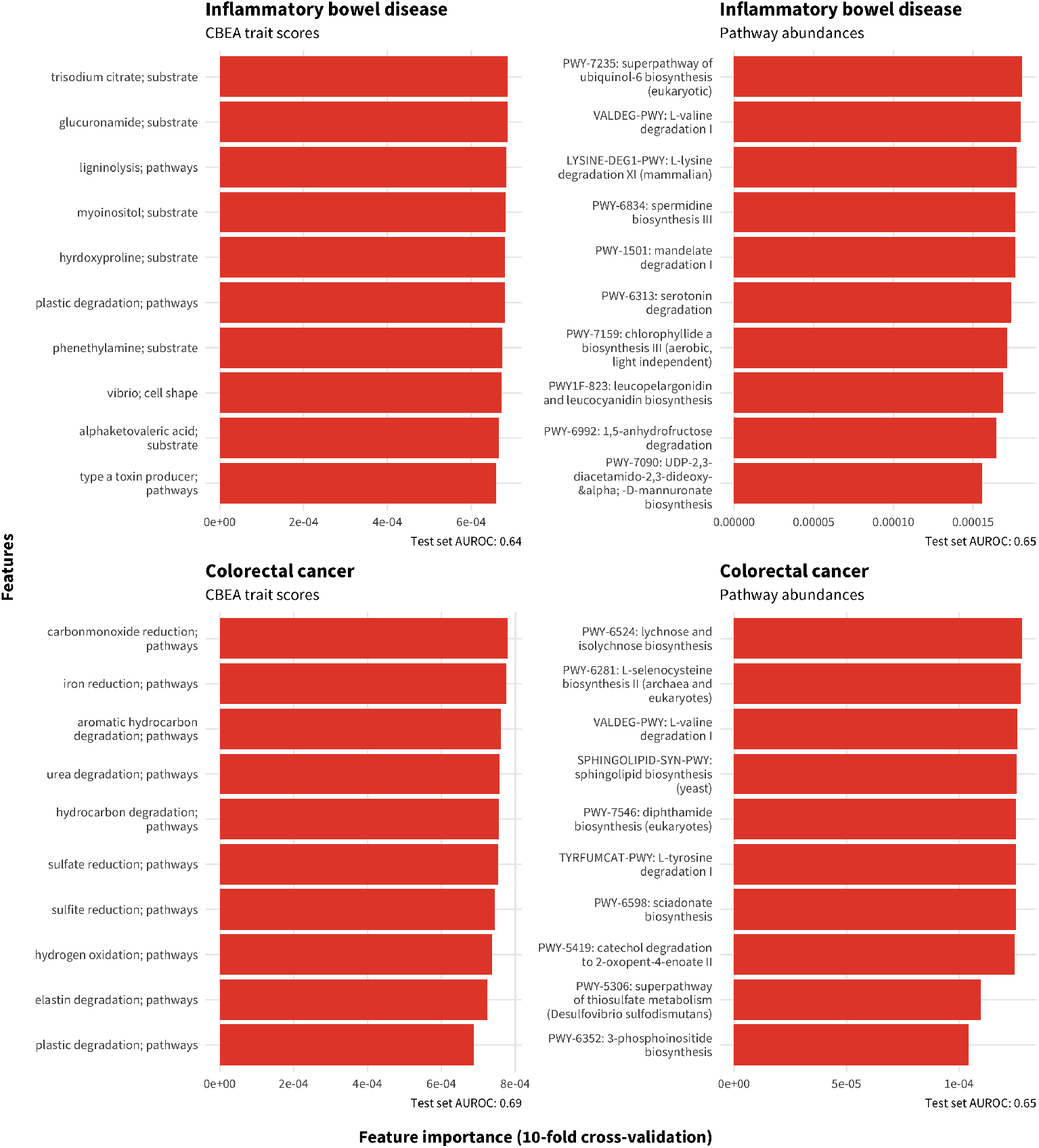
Top 10 important features based on random forest model fitted different inputs from data sets profiled with whole genome metagenomics. Features were selected from mean decrease in Gini impurity averaged across 500 decision trees and 10-fold cross-validation (nested with the training set) as implemented in `scikit-learn`. AUROC scored on a held-out test set is also presented for each input type and disease condition.

## Discussion

### Traits are annotated with high coverage at the species-level

We computed the coverage of trait annotation on a typical dataset to understand the extent in which community function is captured, thereby serving as a proxy for expected confidence for an enrichment analysis performed using trait-based taxon sets. Low coverage in this case indicates that the database does not adequately capture the diversity of microbes found in the target data. This is because there might not be enough taxa present in the data set to serve as evidence for the trait. Alternately, this could also mean that the analysis is missing a majority of underlying community traits, many of which might be core to the health outcome association of interest but simply missing in the analysis. We computed coverage based on two metrics: first, a richness-like metric which computes coverage as the proportion of taxa present per sample annotated to a trait (for a given category and data set); second, an evenness-like metric that accounts for relative abundances of each annotated taxa by computing coverage as the proportion of reads per sample annotated to a trait.

When evaluated on the HMP data set (Fig 1), we can see that the overall richness coverage is low (less than 25% of identified species) across all sites and data sets, particularly for nasal cavity and vaginal sub-sites. However, when considering evenness of coverage, almost all of reads were annotated to a trait. This is consistent with the observation that relative abundances of human-associated microbiomes are highly skewed [16], where a small number of species usually dominate the community. As such, even though traits might only cover a small number of taxa, they might represent the majority of community abundance. For example, Ravel et al. [53] observed that *Lactobacillus* species dominate the vaginal microbiome and, in some phylotypes, almost all reads are assigned to a single species. This shows that our trait-database has high degree of coverage across the most abundant taxon within a community, which supports utilizing these sets to perform exploratory analyses. However, low richness coverage also indicates that our database might not capture traits associated with rare taxa, which can play an important role in regulating host health [58].

Unfortunately, coverage is significantly lower for samples profiled using 16S rRNA gene metabarcoding (Fig S1). For some trait categories, such as pathways, no traits were assigned to any taxa (Fig 2). We hypothesized that this is due to two issues. First, metabarcoding data sets can only resolve taxonomies at the genus level [30], while traits are usually defined at the species and strain levels. Aggregating consensus traits to the genus is difficult due to the high degree of strain and species level diversity within the microbiome [11]. Second, taxonomic assignments for metabarcoding data sets are often based amplification of a specific hyper-variable region for a marker gene (most often the 16S rRNA gene). This means that taxonomic assignment can be sensitive to the choice of region, and can be inaccurate. Furthermore, the choice of taxonomic database (e.g. Ribosomal Database Project [15], SILVA [51]) can also play a part in reducing the ability for trait annotation coverage. Differences between taxonomic paths [1] can result in certain taxa not being able to be matched to traits, which usually assign traits based on NCBI identifiers.

### Trait-set features are predictive of disease outcomes

We assessed the predictive performance of models fit on trait-set enrichment scores compared to other function-based inputs. For whole genome sequencing data sets, measured pathway abundances were utilized as a comparison point while for 16S rRNA gene sequencing data sets, predicted pathway abundances via PICRUSt2 were utilized instead. Fig 2 shows that across all conditions and profiling techniques, trait-set features are competitive in producing well performing models and were able to discriminate between cases and controls. Surprisingly, performance was also comparable in the 16S rRNA gene sequencing data set despite overall low coverage across both richness and evenness metrics. This demonstrates that trait-set abundances can still provide an informative approximation to functional potential similar to PICRUSt2 that can be used for exploratory and hypothesis generating purposes.

To determine which features are important for overall model performance, we extracted the top 10 features based on the mean decrease in Gini impurity. However, the overall feature importance values are not high, suggesting that no individual feature was dominant in classifying patient status. This is further supported by the fact that some nonsense features show up in the top 10 list for models fit using pathway abundances such as PWY-7235 and LYSINE-DEG-PWY, which are mammalian and eukaryotic pathways, respectively. However, the models still show respectable discriminatory power when evaluated on the test set (AUROC ∼ 0.7). Since random forest models can capture interactions between predictors [27], we hypothesized that the interaction between features contribute to test set performance rather than marginal effects. As such, we did not observe a high degree of feature importance scores since these measures are not designed to capture interaction effects [65].

However, despite such limitations, we were still able to recover existing knowledge about the condition of interest. For example, “sulfide reduction;pathways” was shown to be an important feature in discriminating subjects with CRC vs control subjects in Fig 4. This is supported by previous research showing that an increase in abundance of sulfate reducing bacteria is associated with the condition [68]. Mechanistically, this process, when using methionine or cysteine as substrates [13], generates H_2_S as a product, which can stimulate CRC by inhibiting butyrate oxidation (which helps prevent the breakdown of the gut barrier) as well as promoting the generation of reactive oxygen species [42]. Another trait feature is “urea degredation;pathways”, which suggests the importance of bacterial-driven urea hydrolysis process, which is one of the main sources of ammonia in the human gut [5]. Sustained exposure of colonocytes to free ammonia may contribute to the development of CRC [14], which is supported by animal experiments showing histological damage in the distal colon after long-term ammonium exposure [39].

### Limitations and future directions

Even though our results demonstrate that utilizing trait-based sets can provide meaningful insight to microbiome data sets, there are several major challenges to widespread adoption. Although trait databases do not suffer from the same types of biases that exist in genomic reference databases [67], the reliance on curated experimental data means that traits are usually only annotated for species that are well studied and culturable. While using predictive models can help in assigning traits to a broader category of taxa [62], such automated approaches can result in misclassification of traits and increased noise in downstream analyses. Additionally, high-quality trait annotations require a time-consuming, manual curation process [63]. A source that is based on user submission such as GOLD [44] can cover a larger number of taxa and traits, but unfortunately can have erroneous and duplicated assignments due to the lack of a standardized nomenclature. There is currently a gap in producing a high-quality and diverse trait databases that are maintained and continuously updated.

In addition to issues with trait database quality, there are also problems matching the identity of taxa in a given trait database with identifiers found in references for sequence-based taxonomic profiling such as SILVA [51]. For whole genome metagenomic data sets, standard tools (such as MetaPhlan [2]) can provide NCBI identifiers at the species or strain levels. However, it is currently unclear how to aggregate or disaggregate traits if the taxonomic resolution of the observed data set is higher or lower than that of the trait database in use. This is even more difficult with metabarcoding datasets, where low taxonomic resolution makes trait-to-taxa assignments sparse and less confident.

Finally, there are also hurdles in being able to properly validate traits that are found to be significantly enriched due to a lack of ground truth data sets. While some traits can be matched to pathways directly, others involve complex coordination of multiple genetic pathways. As such, further investigation into ways to identify biological concordance between obtained results and external measurements can help improve confidence in utilizing traits for microbiome analyses.

## Conclusion

Set-based enrichment analysis is a useful approach for analyzing microbiome data sets since it not only reflects underlying biology but can also provide more unique perspectives of function that is linked to ecosystem services. Microbial trait databases are a promising resource to construct taxon-sets as traits represent physiological phenotypes. We demonstrated that trait-based sets have high coverage across body sites, especially for samples profiled using whole genome metagenomics. Furthermore, enrichment scores computed on such sets are also competitive in predicting case/control status compared to pathway abundances. As such, trait features found to be important in model fitting can be used to define interesting mechanistic hypotheses.

## Supporting Information

**S1 Fig. Trait coverage statistics for samples profiled with 16S rRNA gene metabarcoding**. Panel **(A)** illustrates the proportion of present taxa per sample annotated to at least one trait. Panel **(B)** illustrates the proportion of reads assigned to taxa annotated to at least one trait which accounts for taxa relative abundances. Each plot facet represents different trait categories that were evaluated. Error bar represents the standard error of the evaluation statistic of interest across the the total number of samples evaluated per body site.

**S2 Fig. Top 10 important features based on random forest model fitted different inputs from data sets profiled with 16S rRNA gene sequencing**. Features were selected from mean decrease in Gini impurity averaged across 500 decision trees and 10-fold cross-validation (nested with the training set) as implemented in `scikit-learn`. AUROC scored on a held-out test set is also presented for each input type and disease condition.

## Acknowledgments

We would like to acknowledge funding provided by National Institutes of Health grants R21CA253408, P20GM130454, R01LM012723 and P30CA023108. The funders had no role in study design, data collection and analysis, decision to publish, or preparation of the manuscript.

